# Mental and Physical Well-being of People Working in Helping Professions, Employed in Youth Educational Centres and Sociotherapy Centres

**DOI:** 10.1101/361311

**Authors:** Justyna Szrajda, Ewa Sygit Kowalkowska, Magdalena Weber-Rajek, Marcin Ziółkowski, Alina Borkowska

**Author notes:** Corresponding author (ESK). **These authors contributed equally to this work.** JSZ and ESK are Joint Senior Authors. **These authors also contributed equally to this work** MWR and MZ and AB.

## Abstract

Representatives of helping professions are particularly exposed to occupational stress. The aim of the study was to evaluate mental and physical well-being, as well as their correlates and predictors in a group of employees working at youth educational centres and sociotherapy centres. A total of 96 employees working at youth educational centres and sociotherapy centres in Kujawsko-Pomorskie voivodeship took part in the study. The following psychometric tools were used: the Psychosocial Working Conditions Questionnaire, the Mini-COPE, the LOT-R, and the GSES. The results obtained indicate that people working in helping professions experience mental and physical health problems. Only 3% of the subjects declared they sleep all night. Over 40% and over 35% of the subjects estimate they suffer from low mood and irritation episodes, respectively, rather frequently or continually. Subjects with poorer mental health are more likely to use Helplessness, Avoidance behaviours, or Turning to religion to cope with stress. The strongest predictor of mental well-being is the sense of self-efficacy. Whereas, the strongest predictor of physical well-being is the ability to cope with stress by giving into the feeling of Helplessness. The study demonstrated poor mental and physical well-being of the subjects. A statistically significant correlation was found between sex and the level of mental and physical health. Employees working at youth educational centres and sociotherapy centres with better mental and physical well-being had a stronger sense of self-efficacy and a higher level of life optimism. Hence, the sense of self-efficacy is a predictor for better mental well-being.

## Introduction

Work conditions are defined as all the factors present in the work environment related to performing work, which affect employees’ health and, if negative, may generate occupational stress [1, 2]. If not eliminated, prolonged occupational stress has a negative impact on employee’s health, deteriorates physical condition, and may cause serious emotional burden [3,4,5]. In the published reports, the occupational group of teachers is described as one of the groups particularly susceptible to stress factors in their work environment [6, 7].

When it comes to employees working at youth educational centres and sociotherapy centres, they are exposed to particular work conditions. The aim of youth educational centres is to eliminate the causes and symptoms of social maladjustment of the youth and prepare them to comply with current social and legal standards. Youth sociotherapy centres are designated for children and youth who due to developmental and social functioning disorders, and learning difficulties require their education and work, as well as sociotherapy methods used, to be individually-adjusted. The most common causes for placing young people in such institutions include: evading compulsory education, signs of demoralisation (use of alcohol, intoxicating substances, and prostitution), aggressive and violent behaviours towards their peers and environment, suicide attempts, running away from home, personality disorders, belonging to an organised criminal group, or committing criminal offences (robbery, vandalism, theft, battery). Therefore, it can be assumed that in addition to the traditional stressors for teachers, the staff in such centres is burdened with additional qualification requirements which are difficult to meet.

Teachers and educators working at such centres belong to helping professions, and are required to work for the well-being of others, show commitment, and establish emotional contact. Occupational load may contribute to mental health problems in people working in helping professions. Due to the particular nature of youth-teacher interactions the staff is exposed to occupational stress. At the same time, the teachers’ educational success depends on their ability to cope with stress [8, 9].

To a substantial extent, the ability to cope with stress factors depends on psychological resources. Personal traits that have a positive effect on health include: self-acceptance, sense of authorship, sense of control, sense of self-efficacy, ability to effectively solve tasks and influence one’s environment, intelligence and flexibility, thoughtfulness and emotional competences, as well as optimism in life [10,11,12].

## The Aim of the Study

The aim of the study was to evaluate mental and physical well-being (interchangeably referred to as ‘mental health’ and ‘physical health’ in this paper), as well as their correlates and predictors in a group of employees working at youth educational centres and sociotherapy centres.

The following study questions were asked:

1. What is the level of mental and physical well-being of the subjects?
2. What is the correlation between the level of the mental and physical well-being, age, job tenure, and sex in the studied group?
3. What is the correlation between the level of mental and physical well-being and personal resources, such as sense of self-efficacy, optimism in life, and strategies for coping with stress in the studied group?
4. Which of the studied personal resources is a predictor for mental and physical well-being in the studied group?

The study was based on scientific reports, therefore we assumed that women will show a lower level of well-being than men, and that dealing with stress, optimism and self-efficacy would be predictors of better health at work. Avoiding behaviours aimed at dealing with stress were a predictor of bad mood [13,14].

## Materials and Methods

The research was approved by the Bioethics Committee of Nicolaus Copernicus University, functioning at Collegium Medicum in Bydgoszcz (KB 628/2016). The study subjects were teachers and educators from five youth educational centres and sociotherapy centres in Kujawsko-Pomorskie voivodeship. 96 subjects (72 women and 24 men), aged 26 to 55 years old (mean age was 42 years ±SD = 9), and with a job tenure of 1 to 30 years (average job tenure was 11 years; ±SD = 6) were enrolled into the study.

The following tools were used in the study:

1. The Psychosocial Working Conditions questionnaire (Polish: “Psychospołeczne warunki pracy”) developed by R. Cieślak and M. Widerszal–Bazyl [15]. The theoretical D scale and two empirical scales (D1 and D2) examining mental and physical well-being were used in the study. The question “What is the level of your well-being?” was the main component of the theoretical scale. The D1 scale included an overall assessment of physical health and stress level, as well as somatic symptoms, such as headache, stomach ailments, and heart problems. The mental well-being factor focused on the evaluation of negative emotional states, life and job satisfaction, and self-confidence. Cronbach’s *α* for the D scale was .94 (.86 for the D1 scale and .83 for the D2). The presentation of the results includes qualitative analyses of the questions, which are acceptable methods for analysing questionnaire results. The collected quantitative and qualitative data were compared with the results for teachers, such as standard ten scores and response frequencies, included in the tool manual.
2. The Mini-COPE (Coping Inventory) is used to measure stress coping abilities. It was developed by Charles S. Carver, and adapted in Poland by Z. Juczyński and N. Ogińska-Bulik [16]. The tool is used to evaluate strategies applied to cope with stress. A subject responds to each statement using a scale from 0 to 3, where 0 means “I almost never do it” and 3 means “I almost always do it”. The higher the result, the more intense is the use of a selected strategy. The factor score structure employs the following strategies: Active coping, Helplessness, Avoidance behaviours, Searching for support, Turning to religion, Acceptance, and Sense of humour. Cronbach’s *α* for the Mini-COPE scale ranged from .79 to .87.
3. The LOT-R (Life Orientation Test) developed by M. F. Scheier, Ch. S. Carver, and M. W. Bridges, and adapted in Poland by R. Poprawa and Z. Juczyński [16]. This test is used to evaluate dispositional optimism understood as a personality trait. Optimism, understood this way, is a generalized positive expectancy for good outcomes. Using a five-grade scale, subjects estimate to what extent a given statement applies to them. Raw results are converted to sten scores in order to evaluate the level of dispositional optimism. Cronbach’s *α* for the entire 6 items of the scale is .78, which suggests that the scale has an acceptable level of internal consistency.
4. The GSES (Generalized Self Efficacy Scale) developed by R. Schwarzer, M. Jerusalem, and Z. Juczyński [16]. The GSES consists of 10 statements which apply to one factor. It measures an individual’s level of perceived self-efficacy in coping with difficult situations and obstacles. The results were then calculated and compared with the standard sten scores. The GSES is intended to test healthy and ill subjects. Cronbach’s alphas ranged from .76 to .90.

Statistical analyses were conducted using the IBM SPSS Statistics 21 package. The Kolmogorov–Smirnov test was used to determine the normality of data distribution. A correlation between variables was evaluated using the Spearman’s *rho* and Pearson’s *r* coefficients. Linear regression analysis with the stepwise method was also used. We assumed a confidence level of *p* < 0.05.

Firstly, the data was analysed using basic descriptive statistics, and the Kolmogorov-Smirnov test was applied to examine the normality of quantitative variable distribution. It turned out that distribution of most variables deviates from the normal distribution. Nevertheless, the skewness value for these variables exceeds the contractual absolute value of 0.8 only in the case of seniority, and consequently the distribution of other scales is not grossly asymmetrical with respect to the Gauss curve. It was therefore decided that parametric tests will be carried out with the participation of other variables.

## Results

The sten scale was used to describe the level of mental and physical well-being of the respondents. The mental well-being results were nearly evenly distributed between three categories. 31 subjects demonstrated poor mental health, whereas 29 and 36 subjects showed average and high mental health, respectively. As far as physical well-being was concerned, 33.3% of the respondents rated it as low, 37.5% as high, and 29.2% as average.

According to the procedure described in the “Proposal for analysis of questionnaire results” chapter in the “Psychosocial work conditions” manual, a percentage of the responses to several questions was used in the analysis. The responses were then compared to the percentage distribution of teachers’ responses in the questionnaire manual (Tab. I, Tab. II).

**Table. I.**
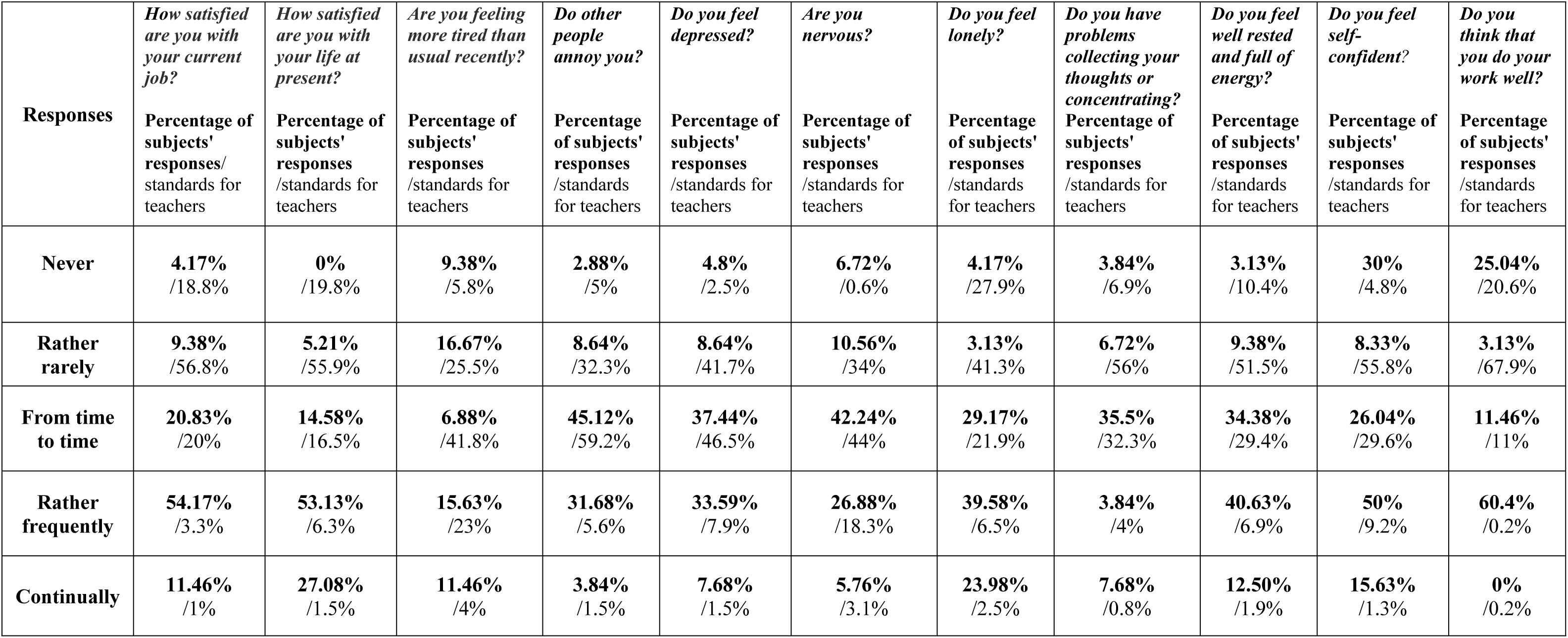
Qualitative analyses of subjects’ responses versus standards for teachers (D2 scale; mental well-being).

| **Responses** | *How satisfied are you with your current job?*<br>Percentage of subjects' responses/ standards for teachers | *How satisfied are you with your life at present?*<br>Percentage of subjects' responses /standards for teachers | *Are you feeling more tired than usual recently?*<br>Percentage of subjects' responses /standards for teachers | *Do other people annoy you?*<br>Percentage of subjects' responses /standards for teachers | *Do you feel depressed?*<br>Percentage of subjects' responses /standards for teachers | *Are you nervous?*<br>Percentage of subjects' responses /standards for teachers | *Do you feel lonely?*<br>Percentage of subjects' responses /standards for teachers | *Do you have problems collecting your thoughts or concentrating?*<br>Percentage of subjects' responses /standards for teachers | *Do you feel well rested and full of energy?*<br>Percentage of subjects' responses /standards for teachers | *Do you feel self-confident?*<br>Percentage of subjects' responses /standards for teachers | *Do you think that you do your work well?*<br>Percentage of subjects' responses /standards for teachers |
| --- | --- | --- | --- | --- | --- | --- | --- | --- | --- | --- | --- |
| **Never** | **4.17%**<br>/18.8% | **0%**<br>/19.8% | **9.38%**<br>/5.8% | **2.88%**<br>/5% | **4.8%**<br>/2.5% | **6.72%**<br>/0.6% | **4.17%**<br>/27.9% | **3.84%**<br>/6.9% | **3.13%**<br>/10.4% | **30%**<br>/4.8% | **25.04%**<br>/20.6% |
| **Rather rarely** | **9.38%**<br>/56.8% | **5.21%**<br>/55.9% | **16.67%**<br>/25.5% | **8.64%**<br>/32.3% | **8.64%**<br>/41.7% | **10.56%**<br>/34% | **3.13%**<br>/41.3% | **6.72%**<br>/56% | **9.38%**<br>/51.5% | **8.33%**<br>/55.8% | **3.13%**<br>/67.9% |
| **From time to time** | **20.83%**<br>/20% | **14.58%**<br>/16.5% | **6.88%**<br>/41.8% | **45.12%**<br>/59.2% | **37.44%**<br>/46.5% | **42.24%**<br>/44% | **29.17%**<br>/21.9% | **35.5%**<br>/32.3% | **34.38%**<br>/29.4% | **26.04%**<br>/29.6% | **11.46%**<br>/11% |
| **Rather frequently** | **54.17%**<br>/3.3% | **53.13%**<br>/6.3% | **15.63%**<br>/23% | **31.68%**<br>/5.6% | **33.59%**<br>/7.9% | **26.88%**<br>/18.3% | **39.58%**<br>/6.5% | **3.84%**<br>/4% | **40.63%**<br>/6.9% | **50%**<br>/9.2% | **60.4%**<br>/0.2% |
| **Continually** | **11.46%**<br>/1% | **27.08%**<br>/1.5% | **11.46%**<br>/4% | **3.84%**<br>/1.5% | **7.68%**<br>/1.5% | **5.76%**<br>/3.1% | **23.98%**<br>/2.5% | **7.68%**<br>/0.8% | **12.50%**<br>/1.9% | **15.63%**<br>/1.3% | **0%**<br>/0.2% |

**Table. II.** Qualitative analyses of subjects’ responses versus standards for teachers (D1 scale; physical well-being).

| **Responses** | *How often do you feel stressed?*<br>(condensed question) | *How would you describe your health as compared to other people at your age?* | *Do you have headaches?* | *Do you often have increased heart rate?* | *Do you experience palpitation?* | *Do you feel weak?* | *Do you feel nauseous?* | *Do you experience chest tightness and/or pain?* | *Do you have stomach problems?* | *Do you fall asleep easily?* | *Do you sleep well?* |
| --- | --- | --- | --- | --- | --- | --- | --- | --- | --- | --- | --- |
|  | Percentage of subjects' responses/ standards for teachers | Percentage of subjects' responses /standards for teachers | Percentage of subjects' responses /standards for teachers | Percentage of subjects' responses /standards for teachers | Percentage of subjects' responses /standards for teachers | Percentage of subjects' responses /standards for teachers | Percentage of subjects' responses /standards for teachers | Percentage of subjects' responses /standards for teachers | Percentage of subjects' responses /standards for teachers | Percentage of subjects' responses /standards for teachers | Percentage of subjects' responses /standards for teachers |
| **Never** | **6.25%**<br>/1.9% | **14.58%**<br>/12.1%<br>(very good health) | **22.91%**<br>/21.3% | **13.54%**<br>/18% | **28.13%**<br>/35.9% | **16.67%**<br>/11.5% | **43.75%**<br>/41.5% | **28.12%**<br>/39% | **26.04%**<br>/34.6% | **27.08%**<br>/26.5%<br>(very easily) | **2.88%**<br>/47.5%<br>(usually I do not wake up at night) |
| **Rather rarely** | **37.5%**<br>/20.6% | **13.54%**<br>/41.5%<br>(rather good health) | **29.16%**<br>/32.1% | **38.54%**<br>/39.7% | **28.13%**<br>/30.5% | **19.80%**<br>/35.6% | **35.42%**<br>/44.8% | **35.42%**<br>/35% | **42.7%**<br>/32.5% | **42.7%**<br>/37.3%<br>(rather easily) | **0%**<br>/30%<br>(wake up once a night) |
| **From time to time** | **33.3%**<br>/43.5% | **33.33%**<br>/39.6%<br>(average health) | **31.25%**<br>/34.2% | **28.13%**<br>/33.1% | **29.17%**<br>/27.3% | **29.17%**<br>/41% | **15.63%**<br>/10.6% | **20.83%**<br>/21.5% | **21.87%**<br>/22.1% | **15.62%**<br>/22.7%<br>(neither easily nor with difficulty) | **18.24%**<br>/17.5%<br>(wake up 2–3 times) |
| **Rather frequently** | **11.45%**<br>/24.6% | **20.83%**<br>/6.3%<br>(rather poor) | **9.37%**<br>/11.7% | **10.41%**<br>/7.3% | **8.38%**<br>/3.8% | **17.7%**<br>/8.3% | **2.08%**<br>/1.9% | **6.25%**<br>/3.5% | **4.16%**<br>/7.7% | **8.33%**<br>/10.8% (with quite difficulty) | **23.04%**<br>/3.5%<br>(wake up 4–5 times) |
| **Continually (very often)** | **11.45%**<br>/9.4% | **17.70%**<br>/0.6%<br>(very poor) | **7.30%**<br>/0.8% | **4.16%**<br>/1.9% | **6.25%**<br>/2.5% | **16.67%**<br>/3.5% | **3.13%**<br>/1.3% | **9.38%**<br>/1% | **5.2%**<br>/3.1% | **6.25%**<br>/2.7%<br>(with big difficulty) | **47.04%**<br>/1.5%<br>(no sleep at all) |

The analyses showed that over 40% and over 35% of the subjects estimate they suffer from low mood and irritation episodes, respectively, continually or frequently (Tab. I). When it comes to sense of nervousness, over 30% of the subjects estimate they “continually” or “rather frequently” experience this state. 40 subjects declared “rather frequent”, and 8 subjects “continual” problems concentrating (which is about 11.52 % of the subjects versus 4.8 % of the norm). More than six times as many respondents declared they experienced loneliness “rather frequently” compared with teachers’ answers presented in the questionnaire manual.

The largest differences between the subjects’ results and the standards for teachers were observed with regard to sleep self-evaluation (Tab. II). Nearly half of the subjects selected the answer “I do not sleep at all”, which is more than six times as many respondents compared with teachers’ answers presented in the manual. Only less than 3% of the subjects declared that they sleep all night – and this result is over sixteen times lower than the official standard. Qualitative analysis also demonstrated that more than five times as many employees working at the centres assessed their health as very bad and quite bad in comparison to the reference group of teachers in the questionnaire manual (38.53% of the sample vs. 6.9% of the teachers). The study participants experienced constant headaches and very frequent tightness and/or pain in the chest more than nine times as often as the reference group of teachers in the questionnaire manual. Nearly 35 % of the respondents reported feeling weak “frequently” or “very frequently” (compared to less than 12 % of the standard quoted in the manual). The respondents indicated that they continually had problems collecting their thoughts or concentrating more than nine times as often as the group of teachers.

Using random samples, a correlation between the level of the mental and physical well-being, age, job tenure, sex, place of work, and job position were evaluated with the Spearman’s *rho* test and Student’s t-test. Correlations between age, job tenure, and the level of mental well-being proved to be statistically insignificant (p = 0.547; p = 0.562). When it comes to the sex variable, the obtained result proved to be statistically significant – *t* (94) = −2.78*;p* < 0.01; 95% *Cl* [-0.73; −0.12]; *d* = 0.664. As the means indicate, women (*M* = 3.34; *SD* = 0.67) scored lower in mental well-being than men (*M* = 3.86; *SD* = 0.50).

The relationship between age, job tenure, and physical well-being also turned out to be statistically insignificant (p = 0.148; p = 0.886). The data obtained using independent samples and the Student’s t-test were statistically significant with respect to the gender variable – t (62.71) = −3.79; p < 0.001; 95% Cl [-0.78; −0.24]; d = 0.811. This means that the level of physical well-being in women (n = 73) is different than that of men (n = 23). As indicated by the mean scores, women (M = 3.59; SD = 0.79) scored lower on this scale than men (M = 4.10; SD = 0.48).

During the next part of the study we analysed the correlation between the level of mental well-being, physical well-being, sense of self-efficacy, optimism in life, and strategies for coping with stress using the Pearson *r* method. The results showed that the level of mental well-being is statistically significantly correlated with strategies for coping with stress, such as Helplessness (r = −0.289; p = 0.004), Avoidance behaviours (r = −0.256; p = 0.012), and Turning to religion (r = −0.224; p = 0.028). These relationships are negative and weak, which means that subjects with poorer mental well-being are more likely to cope with stress by giving into Helplessness, Avoidance behaviours, or Turning to religion. Furthermore, mental well-being is statistically significantly correlated with the sense of self-efficacy (r = 0.401; p = 0.001) and optimism in life (r = 0.291; p = 0.004). These relationships are positive, with the first one being moderately strong, and the second one weak. Which means that the poorer the mental well-being, the lower the sense of self-efficacy and optimism in life.

Physical well-being correlated at a statistically significant level with Helplessness (r = −0.405; p < 0.001), Avoidance (r = −0.222; p = 0.30), Turning to religion (r = −0.315; p = 0.002), sense of self-efficacy (r = 0.376; p < 0.001), and optimism in life (r = 0.346; p = 0.001). Relationships with Helplessness, Avoidance, and Turning to religion are negative, which means that the higher the score for these strategies, the lower the level of respondents’ physical well-being. On the other hand, relationships with the sense of self-efficacy and optimism are positive.

In the final part of the study, we performed a linear regression analysis with the stepwise method. Mental well-being was the dependent variable in the model. The quantitative variables which significantly correlated with this variable were used as predictors. The results of analysis showed that self-efficacy was the strongest predictor, explaining the highest percentage of variance. The *R*^2^ value indicates that about 16.1% of the variability in mental well-being can be explained by the self-efficacy. Furthermore, adding the Avoidance behaviour to predictors’ pool to the model will cause a statistically significant increase in the model efficiency. These two predictors explain 17.9% of the variability of mental well-being. The results are provided in Table III.

**Table. III.**
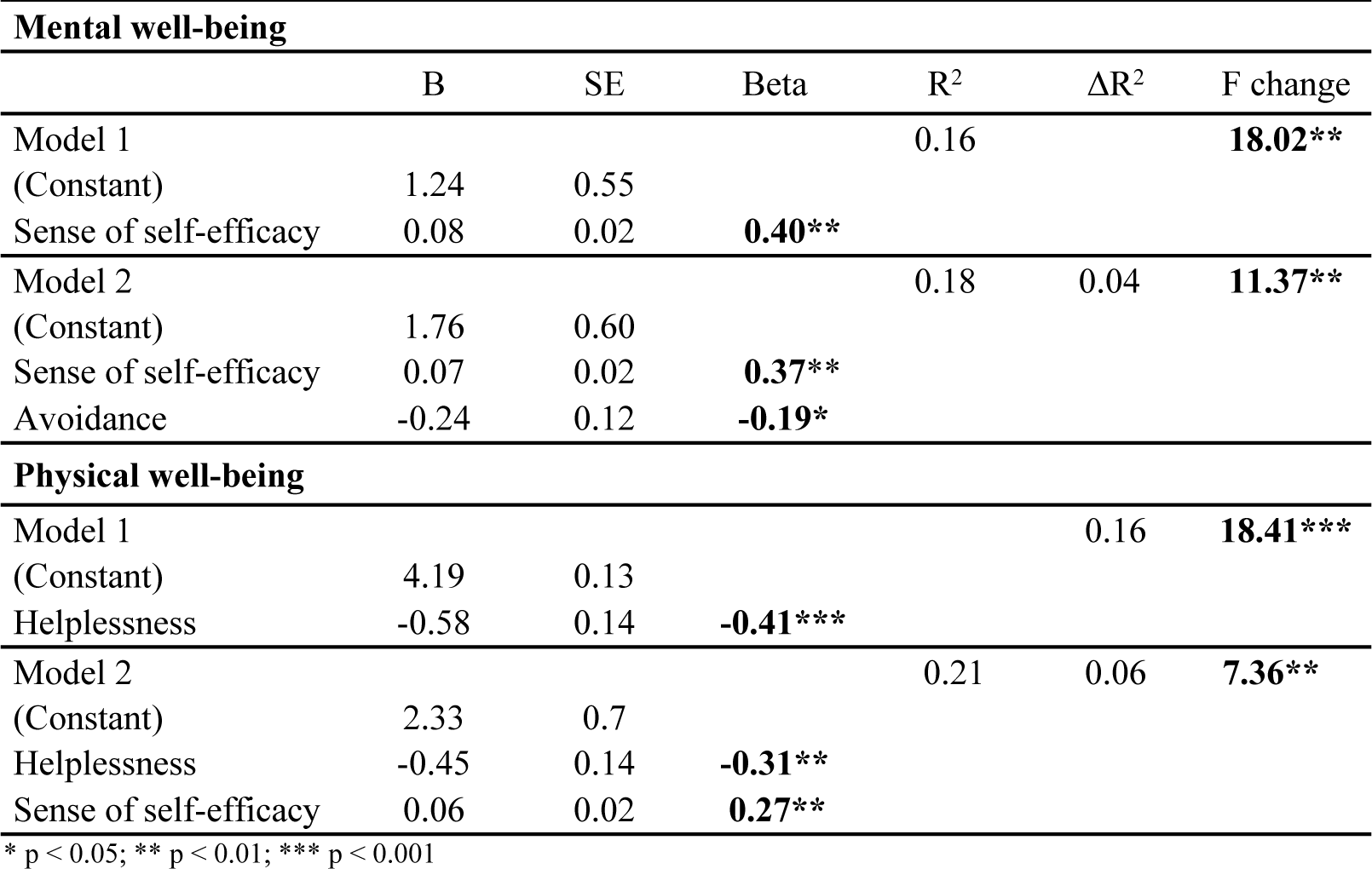
Summary of prediction properties of two models for the dependent variable of mental and physical well-being levels.

The results of analysis showed that Helplessness scale was the strongest predictor, explaining the highest percentage of variance in physical well-being (Table III) allowing to predict 16% of the physical well-being variance. However, adding second parameter to the model: the sense of self-efficacy causes a 21% increase to overall variance. The unstandardised B coefficient indicates that increase of Helplessness score by one unit resulting in 0.45 unit decrease in physical well-being score. On the other hand, increase in self-efficacy score by one unit causes a 0.06 unit decrease in physical well-being score.

## Discussion

The authors believe that due to the restrictive and controlling nature of sociotherapy, employees working at sociotherapy centres need to be competent and have a particular skillset to work with their residents. The psychosocial work conditions of teachers and educators, their health state, and ability to cope with occupational stress have not been the subject of scientific research so far.

The detailed analysis of physical health scores showed that the respondents frequently assessed their health as bad or quite bad, had more headaches, experienced tightness and/or pain in the chest, and more often felt weak and distracted. S. Seidman and J. Zager [17] have already shown that among teachers’ physical ailment symptoms may reflect difficulties in coping with occupational stress and may manifest as pain, depressive disorders, and a tendency to suffer from colds. Somatic symptoms, such as chronic fatigue, migraine headaches, insomnia, and gastric disorders are also typical signs of emotional exhaustion, which may lead to professional burnout [18,19].

The authors believe data on sleep patterns of the respondents should be investigated further. The results of this study are consistent with the data on this professional group available in the literature. The study conducted by C. Reece-Peters among German teachers showed that nearly 30% of the subjects experienced sleep problems [20]. J. C. de Souza et al. [21] showed that secondary school teachers presented symptoms of partial sleep deprivation and poor quality of sleep. This may contribute to excessive sleepiness during the day, affect employees’ health, quality of life, and teaching abilities. Sleep disorders are a very common complication of depression and may be associated with an increased risk of significant depression and suicidal tendencies [22]. Whereas, sleep disorders caused by chronic stress can contribute to the risk of occupational burnout. At the same time, it was found that increased physical activity and proper sleep hygiene ensure high quality of sleep and may buffer the adverse effects of occupational stress [23-25].

In this study, the sense of self-efficacy proved to be the strongest predictor of mental well-being. The authors proved that the sense of self-efficacy is a measure of predicting educational efficacy and ability to solve pedagogical problems [26,27]. The higher the sense of self-efficacy, the higher the objectives people set for themselves and the stronger is their commitment [28]. Teachers with a high sense of self-efficacy are more motivated and efficient in their work. A positive image of oneself as an employee, measured through own actions and efficacy, helps to foster a positive perception of one’s mental health [29].

In this study, sex was a diversifying factor for the study group in terms of the mental and physical well-being. Published literature generally supports the idea that indicators for mental disorders are significantly higher in women than men (33.2% versus 21.7%), also when a professional activity variable is considered (55.4% versus 4.6%) [30,31]. Higher depression and anxiety indicators in the group of women are associated with work factors, such as high requirements and low decision-making power [32]. Studies show that women are particularly exposed to occupational stress factors including multiple roles conflict, discrimination, stereotypes, and a lack of career development [33].

Interestingly, it turned out that work experience, which is another socio-demographic variable, did not influence the mental and physical well-being of examined employees. As other analyses have shown, it can be a significant, positive predictor of job satisfaction in a group of teachers [34, 35]. People working in helping professions are exposed to stress factors which may affect their mental and physical well-being. Due to the particular nature of sociotherapy and the work environment in sociotherapy centres, teachers and educators are required to adapt and act in a specific way to be effective. Therefore, in this study we attempted to evaluate the influence of strategies for coping with occupational load on subjects’ mental and physical well-being. The strategies of Helplessness, Avoidance behaviours, and Turning to religion correlated with mental well-being. These strategies contrast with proactive approach and actively working towards making a positive change. Studies conducted on other helping professions, such as hospital nurses and social workers, indicate that exhibiting passive behaviours in response to stress and lack of support may contribute to emotional exhaustion at work, being a symptom of professional burnout [36, 37].

The correlation analyses showed that mental and physical well-being at work is linked to the level of optimism in life. This is confirmed by studies in secondary school teachers, which show a negative relationship between the optimism in life and causes of occupational burnout, such as personal commitment and emotional exhaustion [38]. Pessimists are more prone to experience physical exhaustion and a lower sense of professional accomplishments.

In this study, 30–40% of the employees suffered from frequent or continual states of irritation, depression, stress, and nervousness. People who have problems coping with occupational stress are more prone to experience the above-mentioned conditions, pain and colds [39]. On the other hand, the ability to recognise, express, and control one’s emotions (i.e., emotional intelligence) influences the ability to cope with stress in helping professions. Also, it may affect the relationship between perceived stress and consequences resulting from certain associated experiences [40,41].

The presented study has its limitations. Workplace stressors were not included in the presented results. For example, we did not analyse the following issues which can play a significant role in regulating stress: dealing with discipline in the classroom, conflicting expectations of teachers and students, and performing tasks under time pressure. The authors of the study did not control any non-work factors, such as other stressful events in a subject’s personal life, that could have potentially influenced teachers’ mental well-being.

According to Ch. Kyriacou [42], studies conducted on a group of teachers should focus on the effectiveness of interventional strategies intended to reduce stress level. Occupational burnout and its influence on teachers’ mental and physical well-being are other aspects that should be studied more thoroughly. The available data indicate that this adverse phenomenon may explain the relation between high expectations at work and poor health of people working in helping professions [43].

## Conclusions

1. People working in helping professions and employed at youth educational centres and sociotherapy centres have difficulty sleeping. Sleep disorders caused by chronic stress may be associated with an increased risk of depression.
2. It was found that women reported significantly worse mental and physical well-being at work than men.
3. People with poorer mental and physical well-being are more likely to cope with stress by giving into Helplessness, Avoidance behaviours or Turning to religion.
4. People with better mental and physical well-being have a stronger sense of self-efficacy and a higher level of optimism in life.

